# A fish frontier? Itatiaia expedition and biodiversity repositories reveal gaps in fish occurrences in Brazil’s high-altitude aquatic ecosystems

**DOI:** 10.1101/2024.11.04.621804

**Authors:** Gustavo Henrique Soares Guedes, Carlos Henrique Pacheco da Luz, Victória de Jesus Souza, Francisco Gerson Araújo

**Author notes:** (corresponding author) (FGA). (GHSG), (VJS), (CHPL).

## Abstract

Brazil harbors one of the greatest diversities of freshwater fish in the world; however, the presence or absence of fish in high-altitude aquatic ecosystems remains largely unknown. This study aims to investigate fish occurrence on the Itatiaia Plateau (Itatiaia National Park) at altitudes ranging from 2,140 to 2,543 meters, marking one of the highest-altitude fish surveys conducted in Brazil. Additionally, it analyzes gaps in fish distribution above 2,000 meters in Brazil by compiling approximately 1 million occurrence records from digital biodiversity repositories (SpeciesLink, Salve, SIBBr). Results from the Itatiaia expedition and biodiversity repositories converge to indicate a lack of fish records in high-altitude aquatic ecosystems (> 2,000 m) in Brazil. *Psalidodon scabripinnis* (Jenyns 1842) is the species with the highest recorded altitude (∼1,944 m). Challenging climatic conditions, physical barriers to dispersal, isolation, historical absence, sampling gaps, and repository biases may be associated with this lack of fish occurrences. This study highlights gaps in knowledge of fish distribution and the potential for future research to discover previously unknown species or species adapted to high altitudes in Brazil.

## INTRODUCTION

By exploring high altitudes and meticulously recording his observations, Alexander von Humboldt (1799–1804) paved the way for understanding the unique patterns and processes that shape biodiversity in mountainous environments (Wulf, 2015). Each altitudinal level harbors complex and diverse ecosystems where life adapts to survive under challenging climatic, spatial, and geomorphological conditions (Fischer et al., 2011; Cassemiro et al., 2023). These vertical landscapes function not only as natural barriers but also as cradles of specialization (Ribeiro, 2006; Buckup et al., 2011; Schluter, Pennell, 2017), where even today, unknown species may still be waiting to be described.

Although Brazil stands out for its vast continental territory and is considered the most biodiverse country in the planet (World Population Review, 2024), its range of mountains and altitudinal gradient is relatively narrow, especially compared to neighboring countries along the Andes mountain range. The Pico da Neblina, at 2,993 meters above sea level, located in the state of Amazonas, represents the country’s highest point (IBGE, 2023). Although advances have expanded our understanding of biodiversity in Brazil (e.g., Boeger et al., 2024), the distribution of fish species at high-altitudes mountains exemplifies the Wallacean shortfall (Hortal et al., 2015) and represents an uncharted frontier. These knowledge gaps are not exclusive to Brazil; studies indicate that fish remain among the least studied taxonomic groups along altitudinal gradients (Fischer et al., 2011). A recent example includes expeditions to the Pico da Neblina National Park, which led to the discovery of new frog species (e.g., Fouquet et al., 2024) and insights into insect community distribution patterns along elevation gradients (Shimabukuro et al., 2023). However, the occurrence of fish on Brazil’s highest mountain remains unknown.

Another significant high-altitude site is Pico das Agulhas Negras in Itatiaia National Park, standing at 2,789 meters, the fifth-highest peak in Brazil (IBGE, 2023). Its relative height, however, is approximately 489 meters, as it rises from the Itatiaia plateau, which is situated around 2,300 meters above sea level (Faria, 2005). This plateau hosts high-altitude fields that develop on the upper terraces of the alkaline massif (Safford et al., 1999), interwoven with an extensive drainage network of streams, temporary marshes, small lakes, and waterfalls. In addition to its impressive elevations and abundant water resources, the plateau sustains a rich biodiversity, including plants, invertebrates, amphibians, reptiles, birds, and mammals adapted to high altitudes (e.g., Ribeiro et al., 2007; Carvajalino-Fernández et al., 2013; Abreu et al., 2017; Aximoff et al., 2020), establishing itself as a complex ecosystem and essential biodiversity refuge in the Serra da Mantiqueira (southeastern Brazil).

To date, fish have only been recorded at lower altitudes in the park (700—1600 m) (Caramaschi, Caramaschi, 1991; Luz et al., 2014), and not on the Itatiaia plateau. The only known expedition referencing fish on the plateau was conducted over 120 years ago (1901—1903) when naturalists Carlos Moreira and Alípio de Miranda Ribeiro explored the area, finding and describing new species but reporting an absence of fish in their findings (Miranda Ribeiro, 1906). Although they noted this possible absence, the survey did not focus on fish, and no specific sampling locations, sampling effort, or capture methods were described. Thus, the presence or absence of fish on the Itatiaia plateau remains uncertain to this day.

Therefore, the central aim of this study is to investigate the occurrence of fish on the Itatiaia Plateau (Itatiaia National Park) at altitudes between 2,140 and 2,543 meters. Three possibilities were expected: 1-Discovery of species that occur at lower altitudes, expanding their distribution to the Itatiaia Plateau; 2-Discovery of new species unknown to science; 3-Absence of fish occurrence. Additionally, fish occurrences at elevations above 2,000 meters in Brazil were assessed using occurrence records from digital biodiversity repositories (SpeciesLink, Salve, SIBBr/GBIF) and digital elevation models. In summary, this study aims not only to expand knowledge on fish distribution in high-altitude ecosystems but also to highlight these underexplored areas as potential sites for significant discoveries. The research seeks to encourage future expeditions that may uncover new species or clarify altitudinal distribution patterns that are still poorly understood in Brazil.

## MATERIAL AND METHODS

### Study area

The Itatiaia National Park (Fig. 1) is located in the Mantiqueira Mountain Range (southeast Brazil; 22°23’02.6” S — 44°40’06.1” W), covering an area of approximately 300 km^2^. The park is dominated by mountainous and rocky elevations, with altitudes ranging from 540 to 2,791 meters, at its highest point, Pico das Agulhas Negras (IBGE, 2023). The area is considered a high-priority site for biodiversity conservation, being the first national park established in Brazil in 1937 and part of the Atlantic Forest Biosphere Reserve (ICMBio, 2013). In the region known as the Itatiaia Plateau, also referred to as the Upper Part, high-altitude grasslands and hanging valleys are present (Safford, 1999), characterized by an extensive network of springs that feed small streams, temporary marshes, lakes, and waterfalls. The waters from this region drain into two main river basins: Paraná and Paraíba do Sul River Basins (ICMBio, 2013).

**FIGURE 1.**
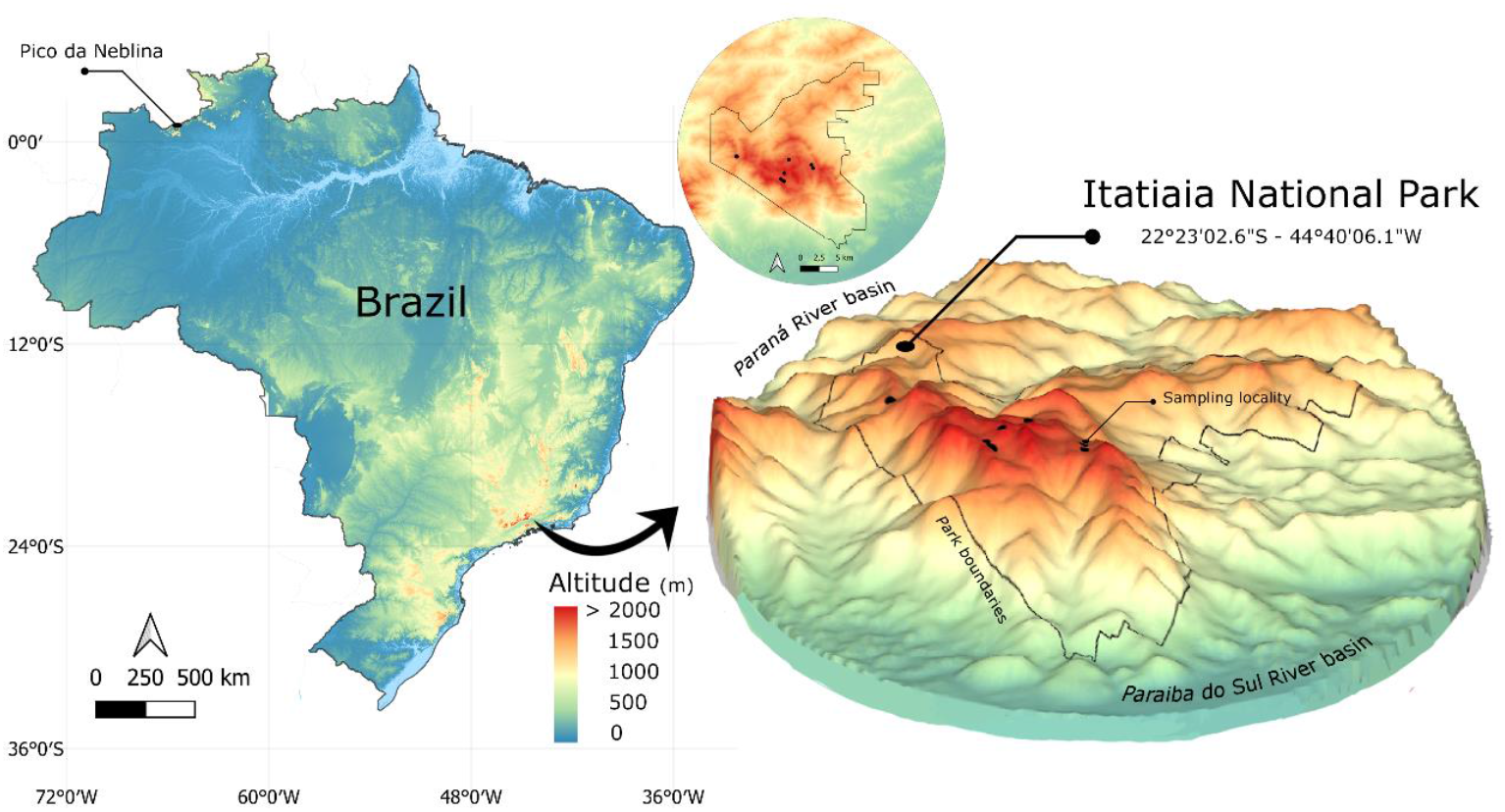
Digital Surface Model of Brazil and Itatiaia National Park. Color scale according to altitude. Black polygon representing the park boundaries. Black points represent sampling localities. Layer: ALOS AW3D30 - Japan Aerospace Exploration Agency.

### Abiotic and biotic sampling

Fish samplings were carried out on the plateau of the Itatiaia National Park (altitude between 2,140 and 2,543 meters above sea level; a.s.l) in October 18–20, 2024 using an oval hand net (50 × 40 cm, 1 mm mesh), covering different environments (small lakes and dams, streams, waterfalls, and swamps; Tab. 1 and Fig. 2). Abiotic parameters of the water were measured using a multi-sensor device (Hanna HI98194 model). The following variables were recorded: water temperature (°C), dissolved oxygen saturation (%), pH, electrical conductivity (mS/cm), turbidity (FTU), and total dissolved solids (g/L).

**TABLE 1.** Geographic coordinates, altitude (meters a.s.l.), and drainage of the sampled locations on the plateau of Itatiaia National Park, southeastern Brazil. Altitude according ALOS AW3D30 - Japan Aerospace Exploration Agency. Geographic coordinate system: WGS84.

| Sample | Locality | Latitude | Longitude | Drainage | Altitude |
| --- | --- | --- | --- | --- | --- |
| 1 | Small lake | 22°22'40.84" S | 44°40'27.15" W | Paraíba do Sul | 2,543 |
| 2 | Aiuruoca Waterfal | 22°21'41.82" S | 44°40'05.27" W | Paraná | 2,373 |
| 3 | Aiuruoca Waterfal | 22°21'41.73" S | 44°40'05.69" W | Paraná | 2,373 |
| 4 | Rancho Caído Waterfal | 22°22'04.17" S | 44°38'19.21" W | Paraíba do Sul | 2,258 |
| 5 | Stream | 22°22'18.77" S | 44°38'11.91" W | Paraíba do Sul | 2,279 |
| 6 | Small dam | 22°23'06.47" S | 44°40'43.34" W | Paraíba do Sul | 2,385 |
| 7 | Flores Waterfal | 22°23'14.73" S | 44°40'31.24" W | Paraíba do Sul | 2,333 |
| 8 | Flores Waterfal | 22°23'16.48" S | 44°40'27.52" W | Paraíba do Sul | 2,312 |
| 9 | Lapa Swamp | 22°21'28.78" S | 44°44'09.52" W | Paraná | 2,143 |
| 10 | Lapa Swamp | 22°21'28.80" S | 44°44'12.60" W | Paraná | 2,138 |
| 11 | Lapa Swamp | 22°21'25.94" S | 44°44'11.35" W | Paraná | 2,140 |

**FIGURE 2.**
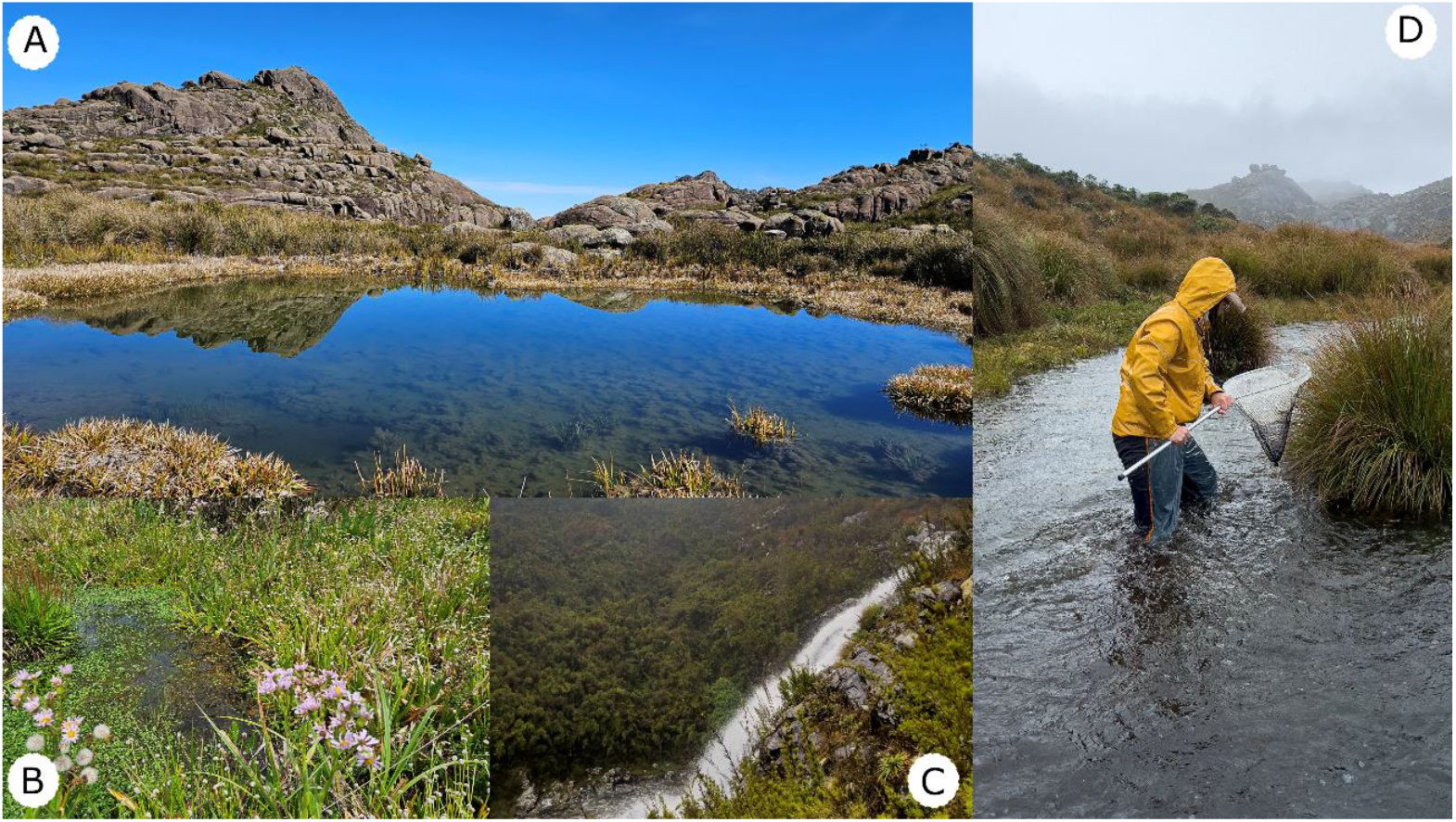
Photographic records of different sampled locations on the plateau of Itatiaia National Park, southeastern Brazil. **A**. small lake; **B**. Lapa Swamp; **C**. Aiuruoca Waterfall downstream; and **D**. upstream views. *Photographs taken by the authors.

### Biodiversity repositories

To identify potential knowledge gaps regarding fish occurrences above 2,000 meters of altitude in Brazil, data were compiled from public digital repositories that aggregate open-access information from museums and biological collections, such as SpeciesLink (specieslink.net), Salve (salve.icmbio.gov.br), and the Brazilian Biodiversity Information System (SiBBr; sibbr.gov.br), which serves as the Brazilian node of the Global Biodiversity Information Facility (GBIF). These repositories were selected for their relevance and scope in aggregating biological data. SpeciesLink and SiBBr compile data from major biological collections in Brazil, while Salve includes records associated with collection permits issued, threatened species, and sampling within Brazilian conservation units (ICMBio, 2024). This latter aspect is particularly important because the highest-altitude locations in Brazil are situated within protected areas.

To download data from these repositories, specific primary filters were applied to suit the distinct characteristics of each search system. In the SpeciesLink repository, the primary filter used was ‘Actinopterygii,’ yielding 426,125 records. In the Salve/ICMBio repository, the main filter was ‘continental fishes,’ resulting in 246,313 records. In SIBBr, the primary filters applied were ‘Actinopterygii,’ ‘present,’ and ‘Brazil,’ with the attribute set to ‘collection,’ yielding 325,445 records. A total of ∼1 million (997,893) occurrences records were extracted from the three repositories. The searches in digital repositories were conducted in October 2024. All records consulted in the repositories are available in the Supplementary Information S1.

To determine the altitude (meters above sea level) of each occurrence record, geographic coordinates were overlaid onto a global Digital Surface Model (DSM) layer set with an approximate horizontal resolution of 30 meters (∼1 arc second), provided by the Japan Aerospace Exploration Agency (www.eorc.jaxa.jp). The overlay and layer sampling were conducted in QGIS software, version 3.32 Lima (QGIS Core Team, 2024), using the ‘Point Sampling Tool’ plugin and based on the WGS 84 geographic coordinate reference system. All occurrence records with coordinates above 2,000 meters were manually verified on a case-by-case basis to assess the compatibility of coordinates with sampling locations as described in the metadata.

## RESULTS

### Itatiaia expedition

The physico and chemical variables of the streams on the plateau of Itatiaia National Park exhibited a neutral pH tendency (mean ± s.d., 7.1 ± 0.2), cold water temperatures (11.4 ± 0.2 °C), and clear water characterized by low turbidity (2.0 ± 0.2 FNU) and minimal dissolved solids (5.5 ± 0.5 mg/L). Additionally, the water exhibited low conductivity (11 ± 1.1 µS/cm) and oxygen saturation levels nearing 80% (± 8.4). Notably, no fish species were recorded at any of the 11 sampled locations on the plateau, situated at altitudes ranging from 2,140 to 2,543 meters a.s.l.

### Biodiversity repositories

Only six (6) occurrences corresponded to fish found at altitudes above 2,000 meters (Tab. 2). However, all six records exhibited inconsistencies and low precision in their geographic coordinates. The four highest-altitude records in the Salve repository (Supplementary File S1) have arbitrary coordinates estimated using the centroid method, representing the geographic center of a set of locations (e.g., midpoint of a protected area). The two subsequent records of *Trichomycterus brunoi* Barbosa & Costa 2010 (catalog number: MBML–4482) were duplicates, appearing in Salve and SpeciesLink. These records also showed low precision, with coordinates indicating a steep rocky escarpment, devoid of apparent water bodies, contradicting the described collection locality (Tab. 2). Therefore, out of the records analyzed, none at altitudes above 2,000 meters could be valid and accurate. *Psalidodon scabripinnis* (Jenyns 1842) is the species with the highest recorded altitude (∼1,944 m) in Brazil according to the biodiversity repositories analyzed (Tab. 2).

**TABLE 2.** Fish occurrence records at high altitudes (meters above sea level) according to biodiversity repositories (Salve, SpeciesLink, and SiBBr), with comments on coordinate precision.

| Source | Species | Latitude | Longitude | Altitude | Comment |
| --- | --- | --- | --- | --- | --- |
| Salve | *Trichomycterus brunoi*<br>Barbosa & Costa 2010 | -20.4584 | -41.7851 | 2,122 | Imprecise coordinate based on centroid method |
| Salve | *Trichomycterus itatiayae*<br>Miranda Ribeiro 1906 | -22.3566 | -44.6837 | 2,111 | Imprecise coordinate based on centroid method |
| Salve | *Characidium vidali*<br>Travassos 1967 | -22.4888 | -43.0597 | 2,098 | Imprecise coordinate based on centroid method |
| Salve | *Parotocinclus maculicauda*<br>(Steindachner 1877) | -22.4888 | -43.0597 | 2,098 | Imprecise coordinate based on centroid method |
| Salve | *Trichomycterus brunoi*<br>Barbosa & Costa 2010 | -20.4201 | -41.828 | 2,073 | Imprecise coordinate, with coordinates indicating a steep rocky escarpment, devoid of apparent water bodies, contradicting the described collection locality. |
| Specieslink | *Trichomycterus brunoi*<br>Barbosa & Costa 2010 | -20.4201 | -41.828 | 2,073 | Duplicate record with the previous one. |
| Specieslink | *Astyanax scabripinnis*<br>Current status: Valid as *Psalidodon scabripinnis* (Jenyns 1842) | -22.7167 | -45.45 | 1,944 | Literature supports species occurrence at altitudes around 1,920 m (e.g, Néo et al., 2000) |

## DISCUSSION

### Itatiaia expedition

The fish sampling conducted during the Itatiaia expedition represents one of the highest-altitude studies (2,140–2,543 meters) in Brazil’s aquatic ecosystems. However, it was not the first to record the absence of fish in the Itatiaia plateau (Miranda Ribeiro, 1906). There are several hypotheses to explain the absence of fish captures at high altitudes in Itatiaia National Park. One possibility is a historical absence, meaning that fish may have never inhabited these ecosystems due to ancient geological and climatic conditions, or they may have once existed but were later extirpated due to landscape evolution processes (e.g., Val et al., 2022). During the Pleistocene, it is speculated that the Itatiaia plateau may have been covered by ice due to extremely cold climatic conditions, and today, Itatiaia represents one of the coldest areas in eastern South America because of its geographic exposure to southern polar fronts (Safford, 1999). The average annual temperature is about 14°C, with colder temperatures sometimes dropping below -10°C during the approximately 56 nights of freezing temperatures per year (Ribeiro et al., 2007). Local flora and fauna species show adaptations to these extreme climatic conditions, such as a dormancy state in the toad *Melanophryniscus moreirae* (Miranda-Ribeiro, 1920) (Carvajalino-Fernández et al., 2013) and the prevalence of the C3 pathway in plant metabolism (Ribeiro et al., 2007).

Additionally, physical barriers to dispersal, such as waterfalls, rapids, and steep, rugged terrain, may prevent fish from migrating to higher altitudes (e.g., May et al., 2017), hindering the arrival and colonization of fish species that occur at lower altitudes (< 1600 m) in Itatiaia National Park (Miranda Ribeiro, 1906; Caramaschi, Caramaschi, 1991; Luz et al., 2024). In general, a decrease in the number of species toward mountain headwater streams is a broad pattern of the river continuum concept (Vannote, 1980). This decline can be attributed to low temperatures and the steep slopes of mountain streams, which act as natural filters for species dispersal, as observed for fish communities in the Iberian Peninsula (Oliveira et al., 2012) and in the Andes-Amazon transition zone (Bogota-Gregory et al., 2024). Other possibilities, such as food scarcity, low habitat complexity, or unfavorable physical and chemical water conditions, are unlikely to be limiting factors. The water bodies in this region present diverse microhabitats (e.g., rapids, pools, rocks, branches), a high diversity of invertebrates (e.g., Macedo et al., 2016; Ribeiro et al., 2019), which provide ample food resources for fish, as well as water with typical characteristics (excluding temperature), such as neutral pH and good oxygenation, conducive to fish development (Tundisi, Tundisi, 2016).

The absence of fish captures on the Itatiaia plateau may also be due to methodological limitations that were not excluded for in this study and require further investigation. Although the sampling covered approximately 50 km of trails and the main water bodies, the region’s extensive hydrological network implies that not all aquatic habitats were sampled. Alternative methods, beyond the hand nets used in this study, such as electrofishing and environmental DNA (eDNA) analysis, could offer promising solutions for fish detection (Xing et al., 2022). However, these methods also have limitations: the low conductivity of the water could reduce the effectiveness of electrofishing, and the high diversity of bird species in the region could introduce confounding factors for eDNA analysis (e.g., fish remains in bird feces, leading to false positives). These challenges underscore the need for further refinement and complementary approaches in future studies.

### Gaps in Fish Occurrences in High-Altitude Aquatic Ecosystems

*Psalidodon scabripinnis* (Jenyns, 1842) is the species with the highest-altitude valid occurrence record (1,944 m a.s.l) found in the analyzed biodiversity repositories. Collected by Charles Darwin and initially described as belonging to the genus *Astyanax* by Jenyns, the species was later repositioned into the genus *Psalidodon* (Terán et al., 2020). This record is located approximately 87 km from the Itatiaia Plateau, in streams of the Serra da Mantiqueira that drain into the Paraíba do Sul River basin (Néo et al., 2000; Castro et al., 2014a). These fish are considered a “species complex,” with some populations isolated for millions of years in different hydrographic basins and separated by hundreds of kilometers (Moreira-Filho et al., 2004). Allopatric populations at different altitudes exhibit variations in morphometric, reproductive, and chromosomal characteristics, including B chromosomes, which may be associated with distinct selective pressures in each altitudinal zone (Néo et al., 2000; Moreira-Filho et al., 2004; Castro et al., 2014a; Silva et al., 2022). Although the population of *P. scabripinnis* represents a notable example of occurrence at high altitude, the question remains: could this be the altitudinal distribution limit for Brazilian freshwater fish?

This study highlights the absence of valid records of fish occurrences above 2000 meters in altitude in Brazil. High altitudes, even with their inherent extreme climatic conditions, do not appear to limit fish distribution in other high-altitude regions globally, such as the Altiplano of the Andes Mountain Range (3,600–4,500 m) (Vila et al., 2007), nor for the Cyprinidae *Herzensteinia microcephalus* (Herzenstein, 1891), recorded on the northern slope of the Tanggula Mountain on the Qinghai-Tibet Plateau in the Himalayas, considered the highest-altitude fish known in the world, reaching 5,350 m above sea level (Zhu et al., 2021). In addition to the historical, biogeographical, and ecological factors previously discussed for the Itatiaia Plateau, the absence of records at high altitudes in Brazil (> 2,000 m) may be attributed to several other factors. First, this particularly suggests a sampling gap in fish data from high-altitude areas in Brazil, characterizing a Linnean shortfall (knowledge gaps in species taxonomy) or a Wallacean shortfall (lack of knowledge about species distributions) (Hortal et al., 2015). This gap may reflect a sampling bias, as fish research in Brazil tends to focus on more accessible, low-altitude regions where ecosystems are easier to sample (e.g., Junqueira et al., 2020; Lima et al., 2021). Second, although Brazil has a vast territorial extent, altitudes above 2,000 meters are restricted to mountainous areas (e.g., Serra dos Órgãos, Mantiqueira, Imeri, Caparaó, Pacaraima), which are proportionally rarer. Consequently, the likelihood of discoveries at high altitudes in Brazil is reduced to specific and limited mountainous regions.

Third, biases related to records, communication, data availability, and inadequate, incomplete, or absent data reporting represent major barriers to understanding species distributions (Oliveira et al., 2016). For instance, research conducted in high-altitude regions may not have fully reported findings, leading to an underestimation of species presence in these areas. Another contributing factor is the use of secondary databases, where species may have been collected but not formally recorded in biological collections or may be housed in collections without public access via open data platforms. Furthermore, imprecise geographical information, such as the lack of exact coordinates, can underestimate the actual altitude of sampling sites, leading to the exclusion of records that, in reality, occurred above 2,000 meters. This set of factors highlights the need for a rigorous review involving collaboration between field researchers, collection curators, museums, and digital repository administrators. The shared goal should be to ensure the availability of accurate and open data, allowing for a more reliable biodiversity analysis.

## CONCLUSIONS

The expedition to the Itatiaia Plateau and biodiversity repositories converge on evidence indicating the absence of fish distribution records above 2,000 meters in Brazil. However, it is important to note that the absence of records does not necessarily indicate the present, past, or future absence of fish at high altitudes. This study highlights knowledge gaps regarding fish distributions and the potential for discovering previously unknown or high-altitude-adapted species. Reporting absences is critical for understanding species distributions, which are often overlooked in biodiversity studies, posing challenges for species modeling and conservation strategies (e.g., Lobo et al., 2010). Consequently, these findings (or the lack thereof) are expected to encourage further research and future expeditions to determine whether Brazil’s high-altitude mountains represent a frontier for fish.

## ACKNOWLEDGEMENTS

This study was funded by Fundo Brasileiro para a Biodiversidade – FUNBIO Conservando o Futuro, and Instituto HUMANIZE (Proc. # 028/2023), Fundação Carlos Chagas Filho de Amparo à Pesquisa do Estado do Rio de Janeiro (E-26/200.897/2021, E-26/210.103/2023), and Conselho Nacional de Desenvolvimento Científico e Tecnológico (CNPq 140.512/2022-5). Special thanks to the management of Itatiaia National Park and to Marcelo Souza Motta for their support with information, accommodation, and logistics.

## AUTHOR CONTRIBUTION

**Gustavo Henrique Soares Guedes:** Conceptualization, Funding acquisition, Resources, Formal analysis, Investigation, Methodology, Visualization, Writing-original draft, Writing-review, Supervision and editing.

**Carlos Henrique Pacheco da Luz:** Investigation, Methodology, Writing-original draft, Writing-review.

**Victória de Jesus Souza:** Investigation, Methodology, Writing-review.

**Francisco Gerson Araújo:** Supervision, Resources, Writing-original draft, Writing-review and editing.

## DATA AVAILABILITY

The data published in this manuscript are available in supplementary information.

## ETHICAL STATEMENT

Fish were collected under permission of Instituto Chico Mendes de Conservação da Biodiversidade (ICMBio/SISBio ##94739). This study was authorized by the Ethics Council of Animal Use (CEUA/ICBS/ UFRRJ), through Permission12.28.01.00.00.00.45 (02/2023).

